# Chemoreceptor pleiotropy facilitates the functional coupling of the synthesis and perception of mating pheromones

**DOI:** 10.1101/124305

**Authors:** Kathleen M. Zelle, Cassondra Vernier, Nicole Leitner, Xitong Liang, Sean Halloran, Jocelyn G. Millar, Yehuda Ben-Shahar

## Abstract

Optimal mating decisions depend on stable signaling systems because any independent changes in either the signal or its perception could carry a fitness cost. However, since the perception and production of specific mating signals are often mediated by different tissues and cell types, the genetic and cellular mechanisms that drive and maintain their coupling on the evolutionary and physiological timescales remain unknown for most animal species. Here, we show that in *Drosophila melanogaster*, sensory perception and synthesis of an inhibitory mating pheromone is regulated by the action of *Gr8a*, a member of the *Gustatory receptor* gene family. Particularly, *Gr8a* acts as a pheromone chemoreceptor in the sensory system of males and females, and, independently regulates pheromone synthesis in the male fat body and oenocytes. These data provide a relatively simple molecular explanation for how genetic coupling allows for the robust and stable flow of social information at the population level.

## INTRODUCTION

The majority of sexually-reproducing animals use intricate mating signaling systems, which rely on the robust physiological coupling between the production and perception of signals since any independent changes in either the signal or the capacity to sense it would carry a fitness cost (Boake, 1991; Brooks et al., 2005; Hoy et al., 1977; Shaw et al., 2011; Shaw and Lesnick, 2009; Steiger et al., 2011; Sweigart, 2010; Symonds and Elgar, 2008; Wyatt, 2014). Previously published theoretical models have hypothesized that possible solutions for maintaining the physiological coupling of signal production and perception are through strong genetic linkage between the genes that regulate mating signal production and those that mediate its perception, or alternatively, via the action of pleiotropic genes that control both processes (Boake, 1991; Butlin and Ritchie, 1989; Shaw et al., 2011; Shaw and Lesnick, 2009). Both mechanisms could provide a robust molecular solution (Chebib and Guillaume, 2019; Kirkpatrick and Hall, 2004; Lande, 1984, 1981, 1980) for the problem of maintaining a stable and reliable population-level social communication system while still retaining the capacity for future signal diversification, as necessitated for speciation (Chebib and Guillaume, 2019; Hoy et al., 1977; Kirkpatrick and Hall, 2004; Lande, 1984, 1981, 1980; Shaw et al., 2011; Shaw and Lesnick, 2009; Wiley et al., 2012).

Although empirical data in support of the role of gene-linkage or pleiotropy in maintaining the physiological coupling between mating signal production and perception are rare, several studies have provided theoretical and experimental support that pleiotropy likely plays an important role in the evolution and maintenance of the physiological coupling between mating-specific signals and the behaviors they elicit at the population level (Chebib and Guillaume, 2019; Hoy et al., 1977; Shaw et al., 2011; Shaw and Lesnick, 2009; Wiley et al., 2012). However, the complex characteristics of mating behaviors, and the signals that mediate them, present a major barrier for identifying the actual pleiotropic genes and molecular pathways that provide genetic coupling between the production and perception of mating signals (Chenoweth and Blows, 2006; Singh and Shaw, 2012). Furthermore, how the perception and production of mating signals remain stably coupled is particularly puzzling for mating pheromones because their perception is mediated by the peripheral sensory nervous system, while their production is often restricted to specialized, non-neuronal pheromone producing cells (Chung and Carroll, 2015; McKinney et al., 2015; Mucignat-Caretta, 2014). Therefore, whether pleiotropy could function in both the production and perception of mating pheromones, a ubiquitous means of signaling across animal mating systems (Mucignat-Caretta, 2014), remained unknown.

Here we show that, in the vinegar fly *Drosophila melanogaster*, some pheromone-driven mating decisions are coupled to the production of specific pheromones via pleiotropic chemoreceptors. Specifically, we demonstrate that *Gr8a*, a member of the *Gustatory receptor* gene family, contributes to the perception of inhibitory mating signals in pheromone-sensing neurons, as well as to the independent production of inhibitory mating pheromones in oenocytes, non-neuronal pheromone-producing cells in the abdomen of the insect. Together, these data provide a relatively simple molecular explanation for how the perception and production of specific mating pheromones are functionally and genetically coupled.

## RESULTS

Similar to other insect species, in *Drosophila*, cuticular hydrocarbons (CHCs), long-chain fatty acids synthesized by the fat body and oenocytes (Billeter et al., 2009; Gutierrez et al., 2007; Krupp et al., 2013; Makki et al., 2014), provide a hydrophobic desiccation barrier for the insect body, and play an important role as pheromones in regulating diverse behaviors, including mating (Blomquist and Bagnères, 2010; Chung and Carroll, 2015; Ferveur, 2005; Howard and Blomquist, 2005; McKinney et al., 2015). Specifically, complex blends of CHCs are often utilized by insect species to communicate sex identity and female mating status, as well as to define the behavioral reproductive boundaries between closely related species (Billeter et al., 2009; Chung et al., 2014; Chung and Carroll, 2015; Coyne et al., 1994; Dweck et al., 2015; Ng et al., 2014; Shirangi et al., 2009; Yew and Chung, 2015). In *Drosophila*, the perception of volatile CHCs is mediated by olfactory sensory neurons located in the antennae and maxillary palps (Benton et al., 2007; Kurtovic et al., 2007; Lebreton et al., 2014; van der Goes van Naters and Carlson, 2007), while less volatile CHCs, whose perception requires physical contact, are sensed by specialized gustatory-like receptor neurons (GRNs) in the appendages (legs and wings) and the proboscis (Koh et al., 2015; Lu et al., 2014, 2012; Thistle et al., 2012; Toda et al., 2012). While some of the genes and pathways that contribute to CHC synthesis in *Drosophila* are known, the molecular identities of most CHC receptors remain unknown. Previous work suggests that the gene *Desat1*, which encodes a fatty acid desaturase enriched in oenocytes, might also independently contribute to CHC perception (Bousquet et al., 2011). However, due to the expression of *Desat1* in central neurons (Billeter et al., 2009) and the broad impact its mutant alleles have on the CHC profiles of both males and females (Labeur et al., 2002), whether *Desat1* directly contributes to mating signal perception remains unresolved.

Consequently, we chose to examine members of the *Gustatory receptor* (*Gr*) gene family as candidates for pleiotropic factors that might contribute directly to both the perception and production of pheromonal mating signals in *Drosophila*. Because several family members have already been implicated in the detection of specific excitatory and inhibitory pheromones (Bray and Amrein, 2003; Hu et al., 2015; Miyamoto and Amrein, 2008; Moon et al., 2009; Watanabe et al., 2011), and the majority of genes that encode family members are already known to be enriched in GRNs (Clyne et al., 2000; Dunipace et al., 2001; Scott et al., 2001; Wang et al., 2004), we reasoned that any pleiotropic *Gr* genes should be expressed in abdominal oenocytes (Billeter et al., 2009) in addition to GRNs. Accordingly, an RT-PCR screen revealed that 24 out of the 60 genes that encode *Gr* family members in the *Drosophila* genome (Clyne et al., 2000; Dunipace et al., 2001; Robertson et al., 2003; Scott et al., 2001) exhibit enriched expression in abdominal tissues (Table 1).

**Table 1.** Candidate *Gr* genes in male and female abdomen. Plus sign indicates presence of PCR product and minus signs indicates PCR product not detected. Genes with no PCR product detected in male or female abdomens not shown.

| **Gene** | **Male** | **Female** |
| --- | --- | --- |
| *Gr2a* | - | + |
| *Gr8a* | + | - |
| *Gr10a* | + | + |
| *Gr21a* | - | + |
| *Gr22a* | + | - |
| *Gr22e* | + | + |
| *Gr36c* | + | - |
| *Gr58c* | + | + |
| *Gr59a* | + | + |
| *Gr59b* | + | + |
| *Gr63a* | + | - |
| *Gr64a* | + | - |
| *Gr64b* | + | + |
| *Gr64c* | + | + |
| *Gr64d* | + | - |
| *Gr66a* | + | + |
| *Gr89a* | + | + |
| *Gr93a* | - | + |
| *Gr93d* | + | + |
| *Gr97a* | + | + |
| *Gr98a* | + | + |
| *Gr98b* | - | + |
| *Gr98c* | + | + |
| *Gr98d* | + | + |

Although several members of the *Gr* receptor family, including *Gr68a, Gr32a, Gr66a, Gr39a*, and *Gr33a*, were previously linked to the sensory perception of mating pheromones (Bray and Amrein, 2003; Lacaille et al., 2009; Miyamoto and Amrein, 2008; Moon et al., 2009; Watanabe et al., 2011), none of these candidate pheromone receptors were identified in our initial screen for *Gr* genes with enriched mRNA expression in abdominal tissues of either males or females (Table 1). However, *Gr8a*, which we identified as having male-specific expression in the abdomen (Table 1), has previously been implicated in chemoreception with enriched expression in abdomens (Park and Kwon, 2011), and was previously shown to specifically contribute to the sensory detection of the non-proteinogenic amino acid L-Canavanine (Lee et al., 2012; Shim et al., 2015). Because our initial expression screen was based on whole-abdomen RNAs, and it was previously reported that *Gr8a* is also expressed in some abdominal neurons (Park and Kwon, 2011), we next used a GAL4 transgenic driver to examine the spatial expression pattern of *Gr8a* in male and female flies. As was previously reported, we found that *Gr8a* is expressed in 14-16 GRNs in the proboscis (Figure 1A) (Lee et al., 2012) as well as in two paired GRNs in the foreleg pretarsus (Figure 1B) of male and female flies. In addition, the *Gr8a*-GAL4 reporter was also highly expressed in abdominal oenocyte-like cells in males but not females (Figure 1C). The male-specific abdominal expression pattern was further supported by qRT-PCR (Figure 1D). Using immunohistochemistry, we then demonstrated that *Gr8a* is co-expressed with the oenocyte marker *Desat1* (Billeter et al., 2009), as well as in *Desat1*-negative cells with fat-body-like morphology (Figure 1E-G). Finally, by using immunohistochemistry on a CRISPR/Cas9 generated GFP tagged allele of the *Gr8a* locus, we demonstrated that the GR8A protein localizes to the fat body (Figure 1H).

**Figure 1.**
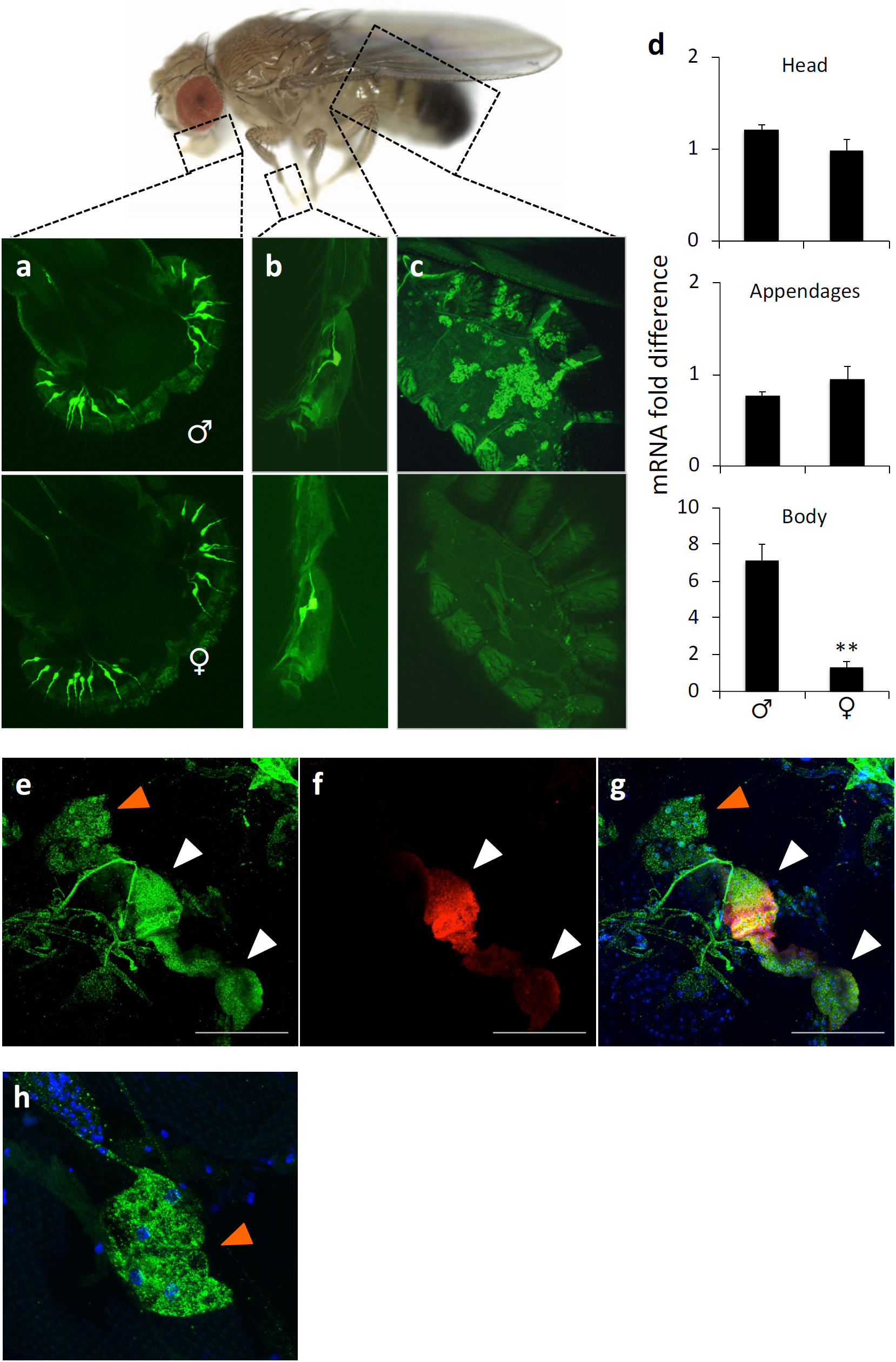
*Gr8a* is a sexually dimorphic chemosensory receptor enriched in male oenocytes and fat body. (A) *Gr8a* promoter activity in proboscis, (B) forelegs, and (C) abdomens of males (top panels) and females (bottom panels). (D) *Gr8a* mRNA expression. Relative mRNA levels were measured by real-time quantitative RT-PCR. **, p<0.01 Mann Whitney Rank Sum Test, n=3/group. (E) Confocal z-stack image of *Gr8a*>EGFP in abdominal cells. (F) Confocal z-stack image of *desat1*>Luciferase in abdominal cells. (G) Co-expression of *Gr8a* and *desat1*. (H) Confocal z-stack image of GFP-tagged GR8A protein in abdominal cells. Green, *Gr8a*; Red, *desat1*; Blue, nuclear DAPI stain. Orange arrowhead, fat body cells; white arrowhead, oenocytes. Scale bar = 100μm.

Together, the enriched expression of *Gr8a* in foreleg GRNs, a primary pheromone chemosensory organ (Lu et al., 2014, 2012), and its sexually dimorphic expression in oenocytes and fat body suggested that in addition to its role in L-Canavanine perception (Lee et al., 2012; Shim et al., 2015), *Gr8a* contributes to mating behaviors. Since mating choices of both male and female flies are determined by a complex blend of excitatory and inhibitory pheromones (Billeter et al., 2009; Krupp et al., 2013), we anticipated that if *Gr8a* is indeed a pleiotropic factor that independently contributes to the perception and production of specific components of the male mating pheromone bouquet, then disruptions of the signal production in males or its perception in females should carry similar impacts on female mate choice behavior. To directly test this hypothesis, we first investigated whether *Gr8a* and the GRNs that express it are required for sensory functions associated with female mate choice by using a single-pair courtship paradigm (Lu et al., 2014, 2012). We found that blocking neuronal transmission in *Gr8a*-expressing GRNs with the transgenic expression of the tetanus toxin (TNT) in females resulted in a shorter copulation latency relative to wild-type females when courted by wild-type males (Figure 2A). Similarly, homozygous (Figure 2B) and hemizygous (Figure 2C) *Gr8a* mutant females exhibited shorter copulation latencies relative to wild-type controls, which could be rescued by the transgenic expression of a *Gr8a* cDNA (Figure 2D). In contrast, similar genetic manipulations of the *Gr8a* gene or *Gr8a*-expressing GRNs in males did not affect courtship behavior as measured by courtship latency and index towards wild-type females (Figure 2-figure supplement 1). These data indicate that *Gr8a* is required for female mate choice via the detection of male-borne inhibitory mating signals.

**Figure 2.**
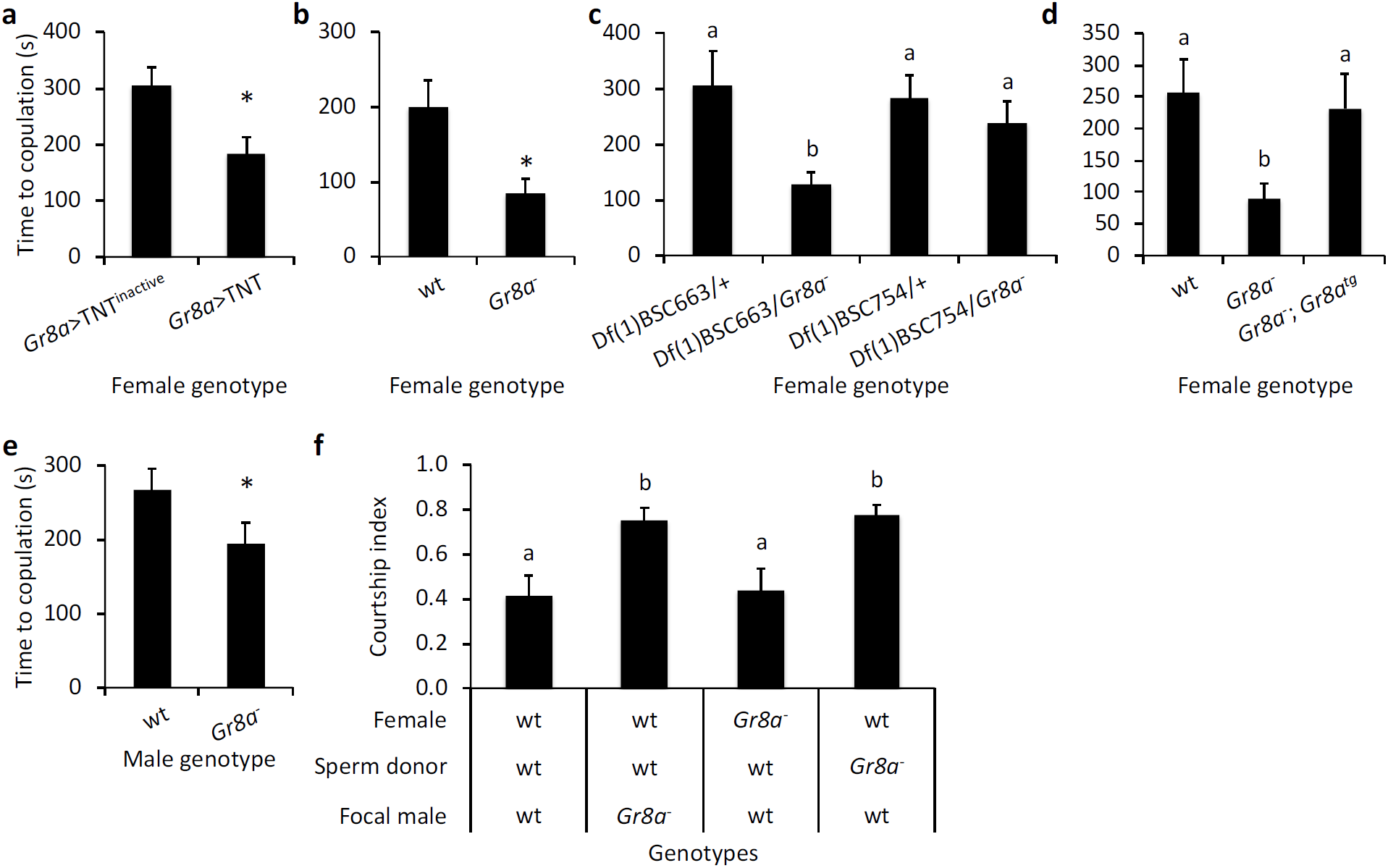
*Gr8a* activity contributes to the perception and production of an inhibitory signal associated with mating decisions in males and females. (A) Blocking neural activity in female *Gr8a*-expressing sensory neurons shortens copulation latency. Homozygous (B) or hemizygous (C) *Gr8a* null females show shortened copulation latency relative to wild-type controls. Df(1)BSC663 is a deficiency that covers the *Gr8a* locus. Df(1)BSC754 was used as a control. (D) Expression of *Gr8a* cDNA with *Gr8a* promoter (Gr8a-;Gr8a^tg^) rescues the copulation latency phenotype in *Gr8a* mutant females. (E) Wild-type females exhibit shorter copulation latency when courted by *Gr8a* mutant relative to wild-type males. (F) *Gr8a* mutant males do not recognize mating status of females, and have a reduced transfer of inhibitory mating pheromones during copulations. Female, female genotype; Sperm donor, genotype of males mated first with focal females; Focal male, genotypes of experimental males presented with mated females. Different letters above bars indicate statistically significant *post hoc* contrasts between groups (panels C, D, and F; p<0.05 ANOVA, n>15/group). *, p<0.05, Mann Whitney Rank Sum Test, n>15/group).

Because *Gr8a* expression is specifically enriched in male oenocytes, we next tested the hypothesis that *Gr8a* also plays a role in the production and/or release of inhibitory mating signals by males. Similar to the effect of the *Gr8a* mutation in females, we found that wild-type virgin females exhibited shorter copulation latencies towards *Gr8a* mutant males relative to wild-type controls (Figure 2E). These data suggest that *Gr8a* mutant males produce and/ or release abnormally low levels of inhibitory mating pheromones.

Mating decisions in *D. melanogaster* rely on a balance between excitatory and inhibitory drives (Kallman et al., 2015). Therefore, in this case, male-borne inhibitory signals may help females optimize mate choice by delaying mounting and sperm transfer from courting males. Additionally, previous studies showed that, in order to increase their fitness, *Drosophila* males transfer inhibitory mating pheromones to females during copulation, which subsequently lowers the overall attractiveness of mated females to other males (Benton et al., 2007; Billeter et al., 2009; Datta et al., 2008; Yew et al., 2009). Because our data indicate that *Gr8a* plays a role in the production of an inhibitory signal in males, we reasoned that the same inhibitory signal may be transferred from males to females during copulation. Therefore, we next tested the hypothesis that *Gr8a* mutant males would have a reduced ability to produce and/or transfer inhibitory pheromones to females during copulation. Accordingly, we found that wild-type males failed to recognize the mating status of wild-type females that were previously mated with *Gr8a* mutant males (Figure 2F). These data indicate that *Gr8a* action is required for the production and/or transfer of inhibitory pheromones from males to females during copulation.

Because *Gr8a* is involved in the perception of an inhibitory mating signal, we also tested the hypothesis that *Gr8a* mutant males exhibit an abnormal capacity for identifying the post-mating status of wild-type females. Indeed, we observed that *Gr8a* mutant males were more likely than wild type males to court a mated female (Figure 2F), suggesting that in addition to its role in the production of inhibitory mating signals, *Gr8a* is also required in males for the sensory recognition of mating inhibitory signals that mark the post-mating status of females. Together, these studies indicate that *Gr8a* is required in males for the production and perception of transferrable inhibitory mating signals that advertise post-mating status in females. These data further support the pleiotropic role of the *Gr8a* gene in the production of inhibitory mating signals in males and the perception of these signals in both males and females in sex-specific contexts.

Because our behavioral data indicate that the *Gr8a* mutation has a dramatic effect on the production and perception of a putative inhibitory mating pheromonal signal, we next examined whether the *Gr8a* mutation has a direct effect on qualitative and quantitative aspects of male and mated-female CHC profiles. Analyses of male CHCs by gas chromatography and mass spectroscopy revealed that the *Gr8a* mutation has a significant impact on the overall qualitative characteristics of the CHC profile of males (Figure 3A), as well as significant quantitative effects on the levels of several specific CHCs in males (Figure 3B-3C and Table 2). In particular, the *Gr8a* mutation affects the levels of several alkenes and methyl-branched alkanes, which have been implicated in mate choice behaviors in diverse *Drosophila* species (Billeter et al., 2009; Chung et al., 2014; Dyer et al., 2014; Shirangi et al., 2009). Similarly, the *Gr8a* mutation affects the expression level of two desaturase enzymes in males, *desat1* and *CG8630* (Figure 3D), which are involved in the biosynthesis of alkenes (Chung and Carroll, 2015). We also found that the overall qualitative aspects of the CHC profiles of wildtype females were not affected by mating with either *Gr8a* mutant or wildtype males (Figure 3E). However, quantitative analyses of individual CHCs revealed that nonacosane (C_29_) is higher in females that mated with the *Gr8a* mutant relative to those that mated with wildtype males (Figure 3F). Together, our behavioral and pheromonal data indicate that *Gr8a* action contributes to mating decisions in females by co-regulating the perception of an inhibitory mating pheromone by females, as well as its production by males, which is consistent with a pleiotropic function for *Gr8a*.

**Table 2.** Male CHCs. Retention time (R.T.), compound, and percent total (% total) of each compound as part of the total pheromonal bouquet for wild-type (wt) and *Gr8a* mutant males.

| R.T. | Compound | wt % total | *Gr8a* % total | p value |
| --- | --- | --- | --- | --- |
| 12.31 | C<sub>21</sub> | 0.589 | 1.102 | <0.001 |
| 13.24 | Unknown | 0.071 | 0.191 | <0.001 |
| 14.2 | C<sub>22</sub> | 0.893 | 0.949 | 0.339 |
| 14.34 | 7-C<sub>22</sub> | 0.362 | 0.490 | <0.001 |
| 15.25 | Unknown | 0.128 | 0.226 | <0.001 |
| 16.22 | C<sub>23</sub> & 9-C<sub>23</sub> | 13.682 | 12.671 | <0.001 |
| 16.4 | 7-C<sub>23</sub> | 30.201 | 35.478 | 0.002 |
| 16.53 | 5-C<sub>23</sub> | 2.772 | 2.389 | <0.001 |
| 16.71 | CvA | 10.240 | 15.700 | <0.001 |
| 18.03 | C<sub>24</sub> & 9-C<sub>24</sub> | 0.578 | 0.367 | <0.001 |
| 18.19 | 8-C<sub>24</sub> | 0.742 | 0.493 | <0.001 |
| 18.27 | 7-C<sub>24</sub> | 0.579 | 0.402 | <0.001 |
| 18.37 | 6-C<sub>24</sub> | 0.395 | 0.233 | <0.001 |
| 18.46 | 5-C<sub>24</sub> | 0.040 | 0.047 | 0.943 |
| 19.09 | 2Me-C<sub>24</sub> | 2.426 | 3.240 | <0.001 |
| 19.95 | C<sub>25</sub> | 0.000 | 2.038 | 0.001 |
| 20.02 | C<sub>25</sub> & 9-C<sub>25</sub> | 6.195 | 1.793 | <0.001 |
| 20.18 | 7-C<sub>25</sub> | 14.781 | 4.344 | <0.001 |
| 20.42 | 5-C<sub>25</sub> | 0.296 | 0.000 | <0.001 |
| 22.89 | 2Me-C<sub>26</sub> | 6.007 | 6.933 | 0.002 |
| 23.7 | C<sub>27</sub> | 1.127 | 0.719 | 0.052 |
| 23.94 | 7-C<sub>27</sub> | 0.308 | 0.083 | <0.001 |
| 26.46 | 2Me-C<sub>28</sub> | 4.992 | 5.775 | 0.078 |
| 27.25 | C<sub>29</sub> | 0.351 | 0.317 | 0.574 |
| 29.89 | 2Me-C<sub>30</sub> | 1.245 | 2.353 | <0.001 |

**Figure 3.**
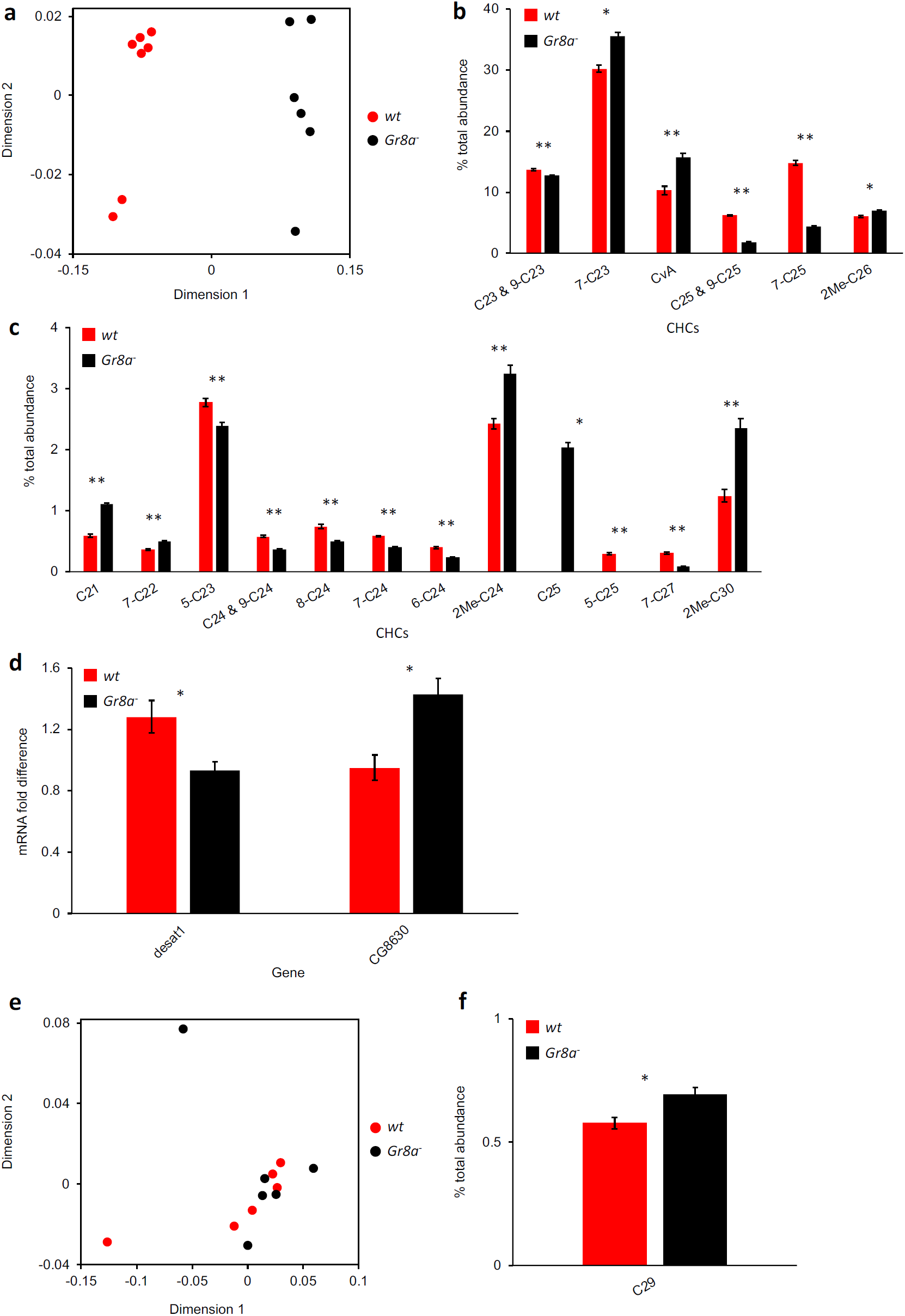
The *Gr8a* mutation affects the pheromone profiles of males and mated females. **(A)** Nonmetric multidimensional scaling (NMDS) plot of CHC profiles of wild-type and *Gr8a* mutant males. p<0.001, permutation MANOVA. **(B-C)** The effect of the *Gr8a* mutation on levels of individual CHCs in males. (B) CHCs found at high proportions in males. (C) CHCs found at low proportions in males. Only affected CHCs are shown. See Table 2 for the complete list. *, p<0.05, **, p<0.001, Student’s t-test or Mann Whitney Rank Sum Test, n=6 (Gr8a-) or 7 (wt). **(D)** The effect of *Gr8a* mutation on the expression level of several desaturase genes. Only affected genes are shown. See Table 3 for the complete list. *, p<0.05, Student’s t-test, n=4/group. **(E)** NMDS plot of CHC profiles of females mated with wild-type or *Gr8a* mutant males. p=0.570, permutation MANOVA. **(F)** Nonacosane (C_29_) differs between females mated with wild-type and *Gr8a* mutant males. See Table 4 for complete list of mated-female CHCs. *, p<0.05, Student’s t-test, n=6/group.

Next, we sought to identify the specific mating inhibitory CHCs that require *Gr8a* for their perception by testing the effect of perfuming males with specific CHCs on wild type female copulation latency in a single-pair courtship paradigm (Lu et al., 2014, 2012). We found that wild-type females did not copulate with *Gr8a* mutant males that were perfumed with the alkenes 9-C_25_, 7-C_25_, and 7-C_27_ (Figure 4). Similarly, we found that wild-type males exhibited a lower courtship index and longer courtship latency when wildtype females were perfumed with 9-C_25_ (Figure 5A-C), and a longer copulation latency when wildtype females were perfumed with 7-C_25_ (Figure 5D-5F). In contrast, perfuming wildtype females with 7-C_27_ had no effect on male courtship or female mating latency (Figure 5G-I). These data suggest that the alkenes 7-C_25_, 9-C_25_, and 7-C_27_ may contribute to an inhibitory signal associated with *Gr8a*.

**Table 3.** Desaturase gene expression. Relative mRNA expression of each desaturase gene for wild-type (wt) and *Gr8a* mutant males.

| Gene | wt mRNA fold difference | *Gr8a* mRNA fold difference | p value |
| --- | --- | --- | --- |
| desat1 | 1.282 | 0.931 | 0.037 |
| desat2 | 1.413 | 1.270 | 0.506 |
| CG8630 | 0.951 | 1.429 | 0.012 |
| CG9747 | 0.838 | 0.525 | 0.343 |
| CG9743 | 1.060 | 0.959 | 0.373 |
| CG15331 | 0.774 | 1.000 | 0.21 |

**Table 4.**
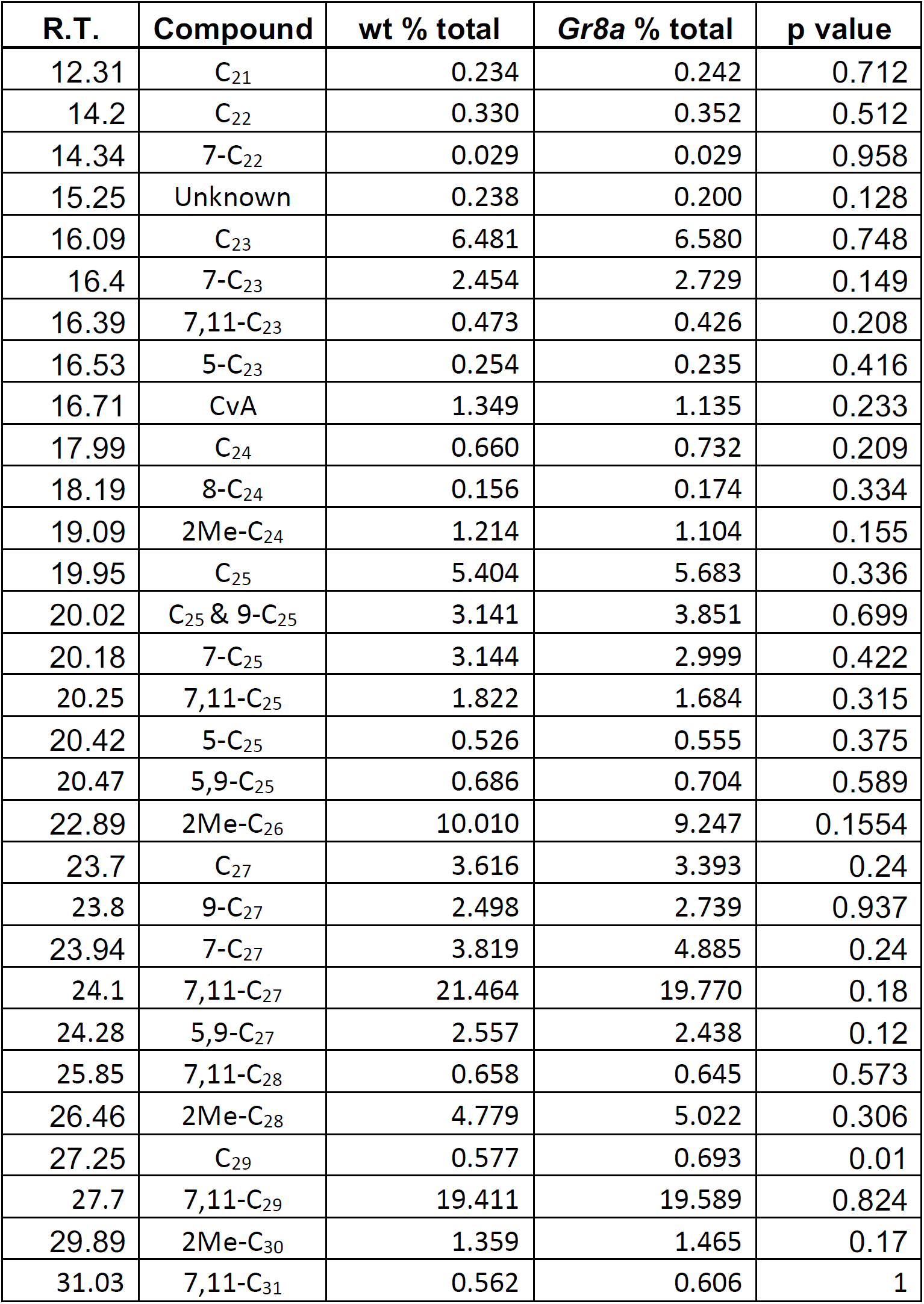
Mated-female CHCs. Retention time (R.T.), compound, and percent total (% total) of each compound as part of the total pheromonal bouquet for females mated with wild-type (wt) or *Gr8a* mutant males.

**Figure 4.**
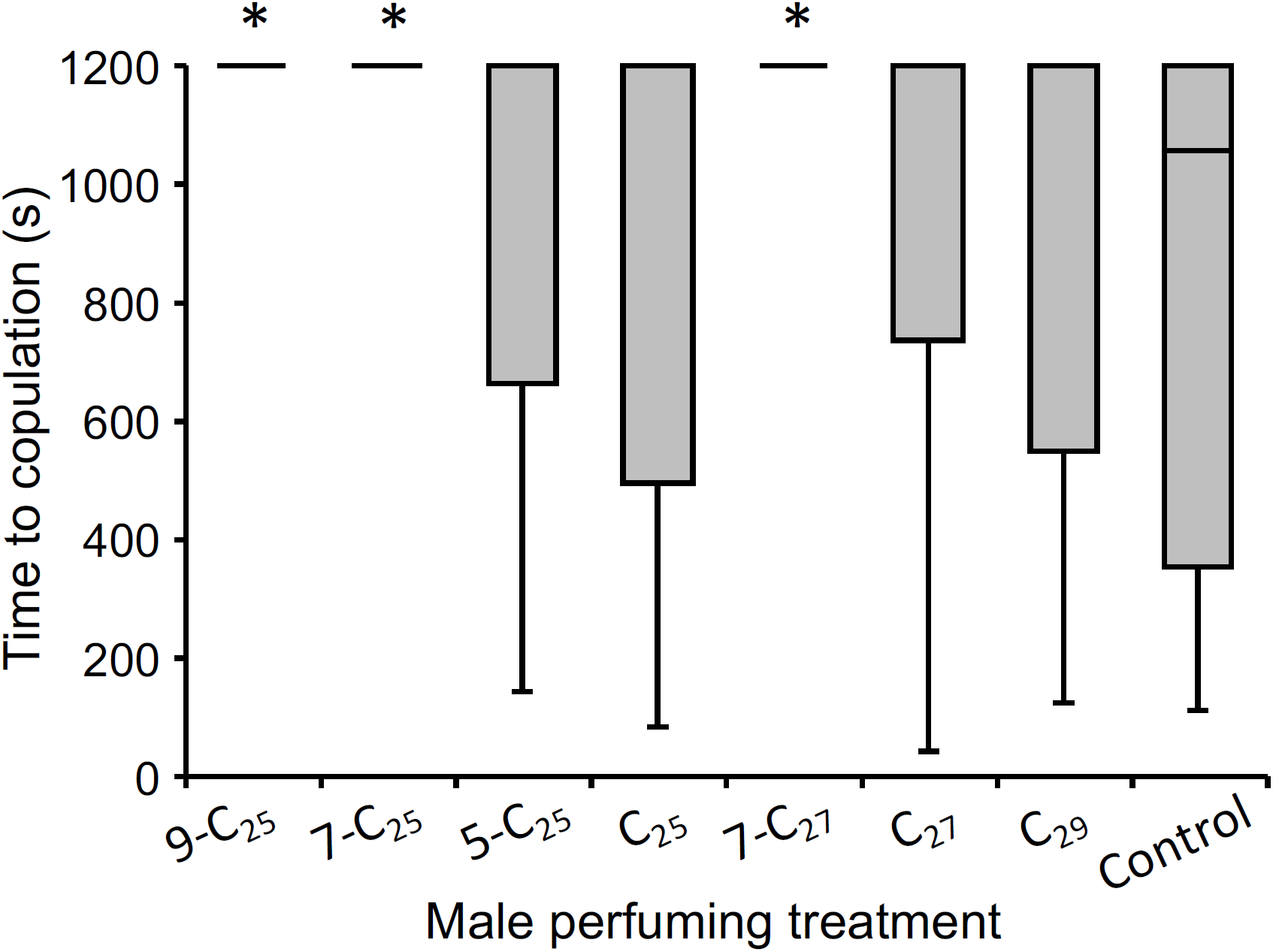
*Gr8a-*associated alkenes inhibit normal courtship behaviors when males are perfumed. Perfuming males with exaggerated amounts of several alkenes increases copulation latency compared to control males. Asterisks above bars indicate statistically significant contrasts compared to control flies, p<0.05, Kruskal-Wallis Test followed by Dunn’s Test, n=15/group.

**Figure 5.**
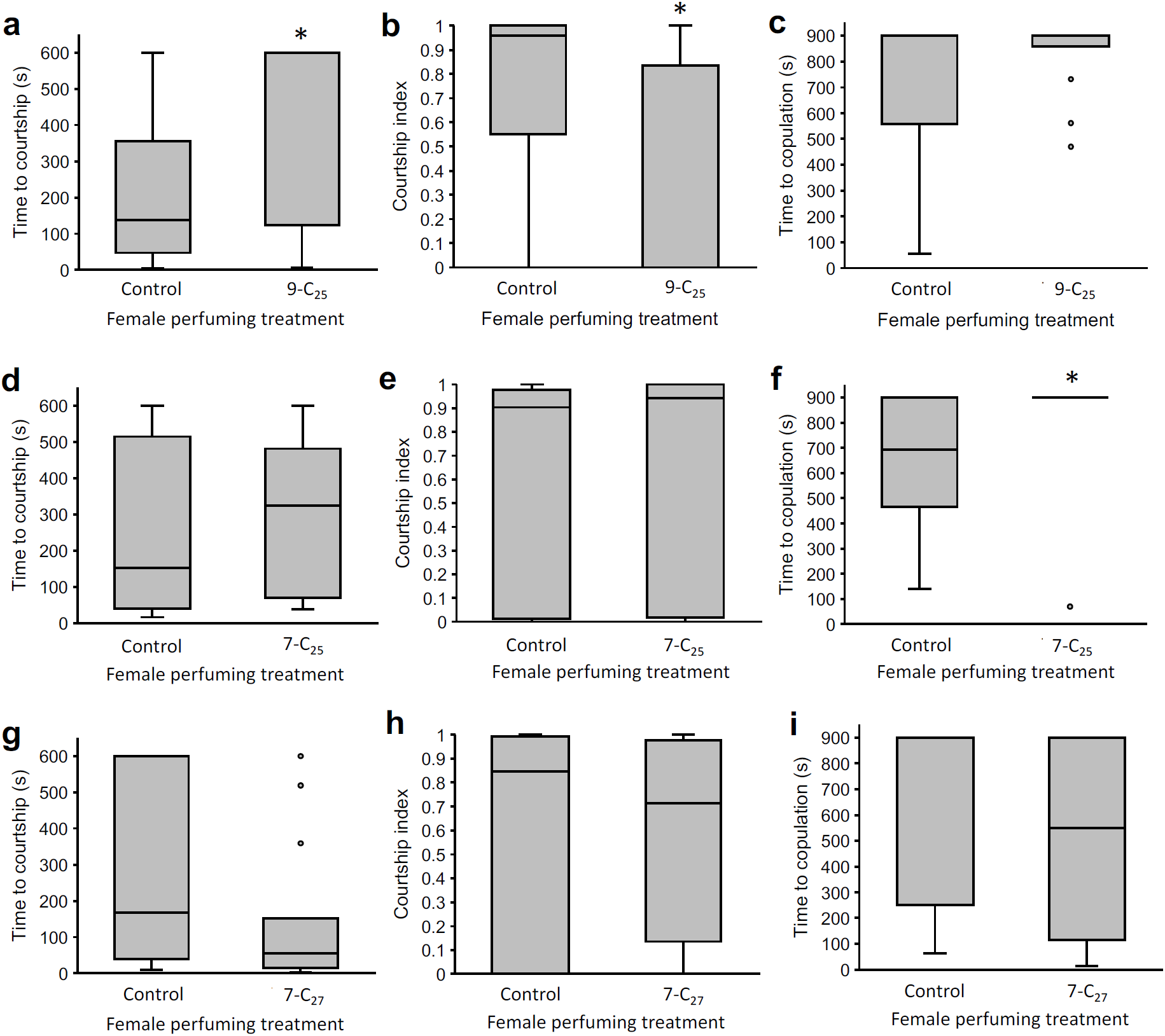
*Gr8a-*associated alkenes inhibit normal courtship behaviors when females are perfumed. Perfuming females with 9-C_25_ increases courtship latency (A), decreases courtship index (B), but does not affect copulation latency compared to control females. Perfuming females with 7-C_25_ does not affect courtship latency (D) or index (E), but increases copulation latency (F) compared to control females. Perfuming females with 7-C_27_ does not affect courtship latency (G), courtship index (H), or copulation latency (I) compared to control females. Asterisks above bars indicate statistically significant contrasts compared to control flies, p<0.05, Mann Whitney Rank Sum Test, n=15/group.

## DISCUSSION

The data presented here demonstrate that *Gr8a* is a pleiotropic chemoreceptor that co-regulates the perception and production of a specific pheromonal signal that plays an important role in mating behaviors of both sexes. In contrast to its expected function in the perception of chemical ligands, how *Gr8a*, a member of a canonical chemoreceptor family might also contribute to the production of pheromonal signals is not obvious. In some better understood secretory cell types, autoreceptors are essential for the regulation of synthesis and secretion rates. For example, dopaminergic and serotonergic cells regulate rates of synthesis and release of their respective neuromodulators by the action of autoreceptors, which regulate synthesis rates via signaling feedbacks in response to changes in the extracellular concentrations of the secreted molecule (Ford, 2014; Stagkourakis et al., 2016). Therefore, one possible explanation for how *Gr8a* might regulate the synthesis and/or secretion of specific CHCs is by acting as an oenocyte-intrinsic autoreceptor, which regulates CHC synthesis rates by providing feedback information about extracellular levels of specific compounds (Figure 6).

**Figure 6.**
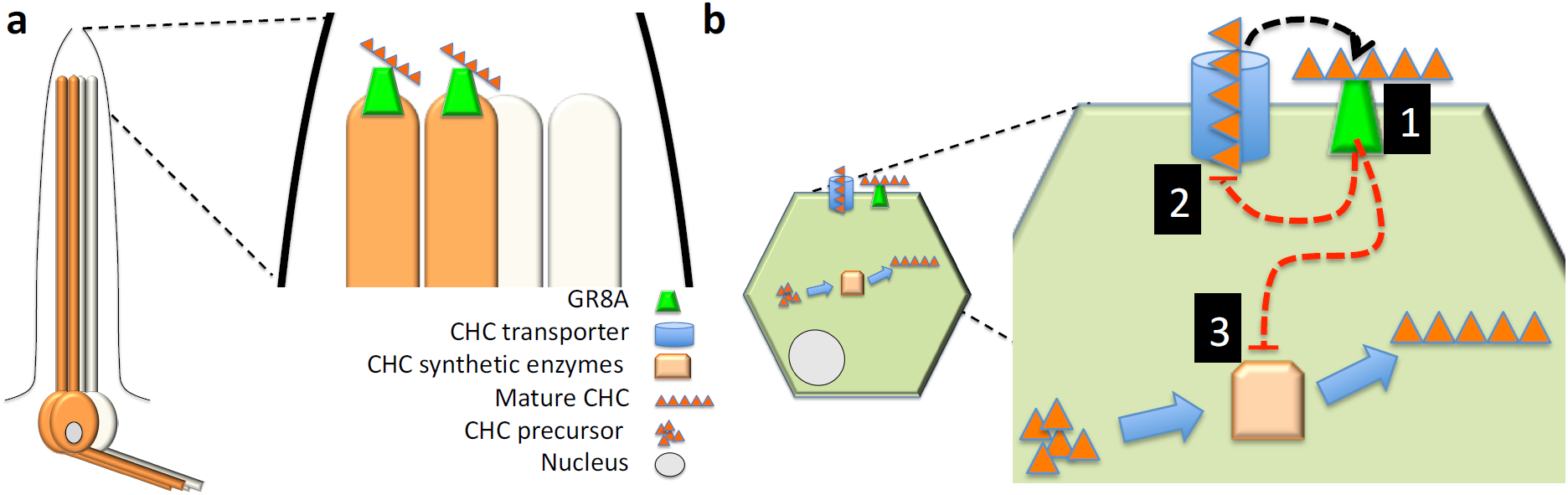
Model for the pleiotropic action of *Gr8a* in the perception and production of pheromones. (A) *Gr8a* functions as a chemoreceptor for an inhibitory signal in pheromone-sensing GRNs of males and females. (B) *Gr8a* also functions as a CHC autoreceptor in oenocytes, which regulates CHC secretion [1] or CHC synthesis [2] via signaling feedback loops [3].

Previous work indicates that *Gr8a* is expressed in “bitter” taste neurons in the proboscis and are specifically required for the sensory perception of the feeding deterrent L-canavanine (Lee et al., 2012; Shim et al., 2015), but not for the detection of other inhibitory chemicals such as caffeine, strychnine, and umbelliferone (Lee et al., 2009; Poudel et al., 2015). Our data indicate that *Gr8a* contributes to inhibitory functions associated with mating decisions as well, as was previously reported for other ‘bitter’ receptors in *Drosophila* (Lacaille et al., 2007; Moon et al., 2009). Together this indicates that ‘bitter’ taste perception and the perception of inhibitory mating signals may share similar pathways in this species.

*Gr8a*-dependent perception of L-canavanine seems to depend on its heterotrimeric interaction with *Gr66a* and *Gr98b* in bitter sensing neurons in the proboscis (Shim et al., 2015). Although both *Gr66a* and *Gr98b* were also identified in our initial screen for receptors enriched in the abdomen, *Gr66a* was expressed in both males and females while *Gr98b* was specifically enriched in females (Table 1). These data suggest that *Gr8a*-dependent contributions to feeding and mating decisions are independently mediated via interactions with different *Gr* genes in each sensory context, and further point to combinatorial subunit composition as an essential aspect of the mechanism for ligand specificity in insect gustatory receptors.

Although we do not yet know the specific chemical identity of the ligand of *Gr8a*, previous studies indicated that at least two inhibitory mating pheromones, 11-cis-vaccenyl acetate (cVA) and CH503, are transferred from males to females during copulations. While our data suggest that the *Gr8a* mutation affects the level of cVA expressed by males, it is unlikely that either cVA or CH503 are the putative *Gr8a* ligands because the volatile cVA acts primarily via the olfactory receptor *Or67d* (Benton et al., 2007; Datta et al., 2008; Kurtovic et al., 2007), and CH503 has been reported to signal via *Gr68a*-expressing neurons, which are anatomically distinct from the *Gr8a* GRNs we describe here (Figure 1A-B) (Shankar et al., 2015; Yew et al., 2009). Instead, our CHC analyses in *Gr8a* mutants (Figure 3), and the results of perfuming behavioral studies (Figures 4-5), suggest that the alkenes 5-C_25_, 7-C_25_, and 7-C_27_ act as inhibitory mating signals as well.

Although the studies we present here do not directly address the possible contribution of *Gr8a* to the rapid evolution of mating signals across *Drosophila*, phylogenetic analyses of *Gr8a* orthologs indicate that this is a conserved, sexually dimorphic receptor across *Drosophila* species (Figure 7A-B). Furthermore, analysis of the *Gr8a* protein alignment revealed that in spite of the high overall sequence conservation, the *Gr8a* protein includes at least one phylogenetically variable domain (Figure 7C-D). These data suggest that in addition to the role of pleiotropic genes, such as *Gr8a*, in maintaining the physiological coupling of the production and perception of mating pheromones, sequence diversification of such genes might also drive the divergence of mating signaling systems, such as the rapidly evolving pheromonal systems across the *Drosophila* species group.

**Figure 7.**
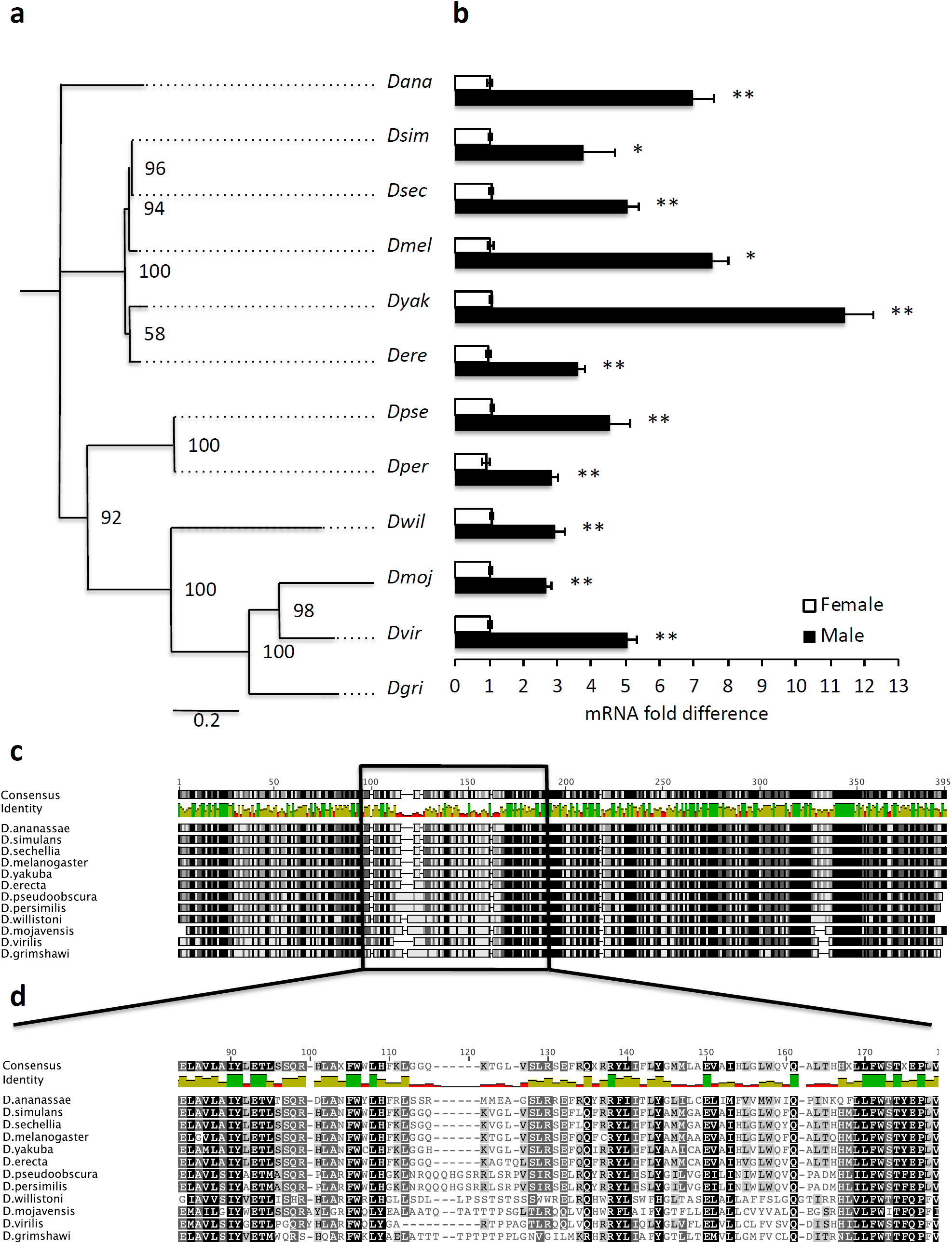
*Gr8a* expression is sexually dimorphic across the *Drosophila* genus. (A) Phylogenetic tree of *Drosophila Gr8a* proteins. Substitution rate = 0.2. (B) *Gr8a* mRNA expression is enriched in males relative to females across *Drosophila*. Black, males; white, females. *, p<0.05; **,p<0.01; Mann Whitney Rank Sum Test, n=4/group. Live *D. grimshawi* was not analyzed because live specimens were not available at the *Drosophila* Species Stock Center (DSSC). (C) Multiple aligned amino acid sequences of *Gr8a* protein sequences from 12 species across *Drosophila.* Box highlights a putative hypervariable protein domain, which is shown at a higher resolution in (D). Numbers on top of alignment indicate amino acid number. Black, 100% identical; Dark Gray, 80-100% similar; Light Gray, 60-80% similar; White, less than 60% similar (Blosum62 score matrix, threshold=1). Bars below consensus represent overall level of amino acid conservation.

Studies in other animal species suggest that receptor pleiotropy likely plays a role in mating signaling via other sensory modalities including auditory communication in crickets (Hoy et al., 1977; Wiley et al., 2012) and visual communication in fish (Fukamachi et al., 2009). While the specific genes and signaling pathways that mediate mating systems in these species are mostly unknown, these data suggest that the genetic coupling of signal-receptor pairs in mating communication systems might be more common than previously thought. Therefore, the genetic tractability of *D. melanogaster*, in combination with the diversity of mating communication systems in this species-rich phylogenetic group, provide a unique opportunity for understanding the evolution and mechanisms that drive and maintain mating communication systems at the genetic, molecular, and cellular levels.

## MATERIALS AND METHODS

### Animals

Flies were maintained on a standard cornmeal medium under a 12:12 light-dark cycle at 25 Celsius. Unless specifically stated, the *D. melanogaster Canton-S (CS)* strain served as wild-type control animals. UAS-mCD8::GFP, UAS-myr::GFP, UAS-TNT-E, UAS-TNT-IMP-V1-A, *Gr8a-GAL4* and *Gr8a*^*1*^ fly lines were from the Bloomington Stock center. Originally in the *w*^*1118*^ background, the *Gr8a*^*1*^ *null* allele was outcrossed for six generations into the *CS* wild-type background. *PromE(800)-GAL4* and *PromE(800)>Luciferase* were from Joel Levine at the University of Toronto. The following *Drosophila* species were obtained from the San Diego Stock Center: *D. simulans* 14011-0251.192, *D. sechellia* 14021-0248.03, *D. yakuba* 14021-0261.01, *D. erecta* 14021-0224.00, *D. ananassae* 14024-0371.16, *D. pseudoobscura* 14011-0121.104, D. *persimilis* 14011-0111.50, *D. willistoni* 14030-0811.35, *D. mojavensis* 15081-1352.23, and *D. virilis* 15010-1051.118. The UAS-*Gr8a* transgenic line was generated by cloning the *Gr8a* cDNA in the pUAST-attB vector using 5’ EcoRI and 3’ NotI restriction sites, followed by *ΦC31* integrase-dependent transgene integration at a Chromosome 2 *att*P landing site (2L:1476459) as previously described (Zheng et al., 2014). For genetic rescue experiments, the UAS-*Gr8a* and *Gr8a*-GAL4 lines were transgressed into the *Gr8a*^*1*^ CS genetic background.

The GFP-tagged allele of *Gr8a* was generated via CRISPR/*Cas9*-dependent editing using a modified “scarless” strategy (www.flyCRISPR.molbio.wisc.edu) as described in (Hill et al., 2019) using the sgRNA CGAGCAAGGCGGGAACGATT, and a donor plasmid containing 1) A backbone with ampicillin resistance; 2) Eye-specific dsRed reporter driven by the 3XP3 promoter, flanked by two *PiggyBac* transposase recognition sites; 3-4) The left and right homology arms, which consisted of 1kb genomic DNA fragments upstream and downstream of the sgRNA site respectively. Single base pair mutations were introduced into the PAM sequence at each sgRNA binding site on the homology arms to prevent Cas9-dependent cutting of the donor plasmid. The left homology arm included the GFP coding sequence in the last intracellular loop of the Gr8a protein. pDCC6 plasmids containing sgRNA and cas9 sequences (100 ng/µL) and the donor plasmid (500 ng/µL) were co-injected into eggs of the *y*^-^*w*^-^background. Subsequently, correct genomic integration of the GFP tag was verified by screening for DsRed-positive animals, followed by sequencing of genomic PCR fragments. Control lines with the same genetic background were generated by selecting DsRed-negative animals. The final tagged *Gr8a* allele was generated by removing the DsRed cassette via the introduction of the *piggyBac* transposase (Bloomington #8285).

### Immunohistochemistry

To visualize the expression pattern of *Gr8a* in males and females, *Gr8a*-*GAL4* flies (Lee et al., 2012) were crossed to UAS-CD8::EGFP and live-imaged at 5 days old using a Nikon-A1 confocal microscope. To demonstrate *Gr8a* expression in oenocytes, abdomens from *Gr8a-*GAL4*/*UAS*-myr::GFP; PromE(800)>Luciferase* flies were dissected and immunostained as previously described (Lu et al., 2012; Zheng et al., 2014) by using a Rabbit anti-GFP (1:1000; A-11122, Thermo Fisher Scientific) and a mouse anti-luciferase (1:100; 35-6700, Thermo Fisher Scientific) antibodies followed by AlexaFluor 488 anti-rabbit and AlexaFluor 568 anti-mouse secondary antibodies (Both at 1:1000; Thermo Fisher Scientific). To visualize the GR8A protein, abdomens of control flies and flies with CRISPR/Cas9 generated GFP-tagged GR8A were dissected and immunostained as previously described (Lu et al., 2012; Zheng et al., 2014) using a Rabbit anti-GFP antibody (1:1000; A-11122, Thermo Fisher Scientific) followed by AlexaFluor 488 anti-rabbit secondary antibody (1:1000; Thermo Fisher Scientific).

### mRNA expression

Newly eclosed flies were separated by sex under CO_2_ and aged for 5 days on standard cornmeal medium. On day 6, flies were placed in a -80°C freezer until RNA extraction. To separate body parts, frozen flies were placed in 1.5ml microcentrifuge tubes, dipped in liquid nitrogen, and then vortexed repeatedly until heads, appendages, and bodies were clearly separated. Total RNA was extracted using the Trizol Reagent (Thermo Fisher Scientific) separately from heads, bodies, and appendages for *Gr8a* expression and from bodies for desaturase enzyme genes. cDNAs were synthesized using SuperScript II reverse transcriptase (Thermo Fisher Scientific) with 500 ng total RNA in a 20 uL reaction. Real-time quantitative RT-PCR was carried out as previously described with *Rp49* as the loading control gene (Lu et al., 2014, 2012; Zheng et al., 2014). Primer sequences are described in Tables 5, 6 and 7.

**Table 5.**
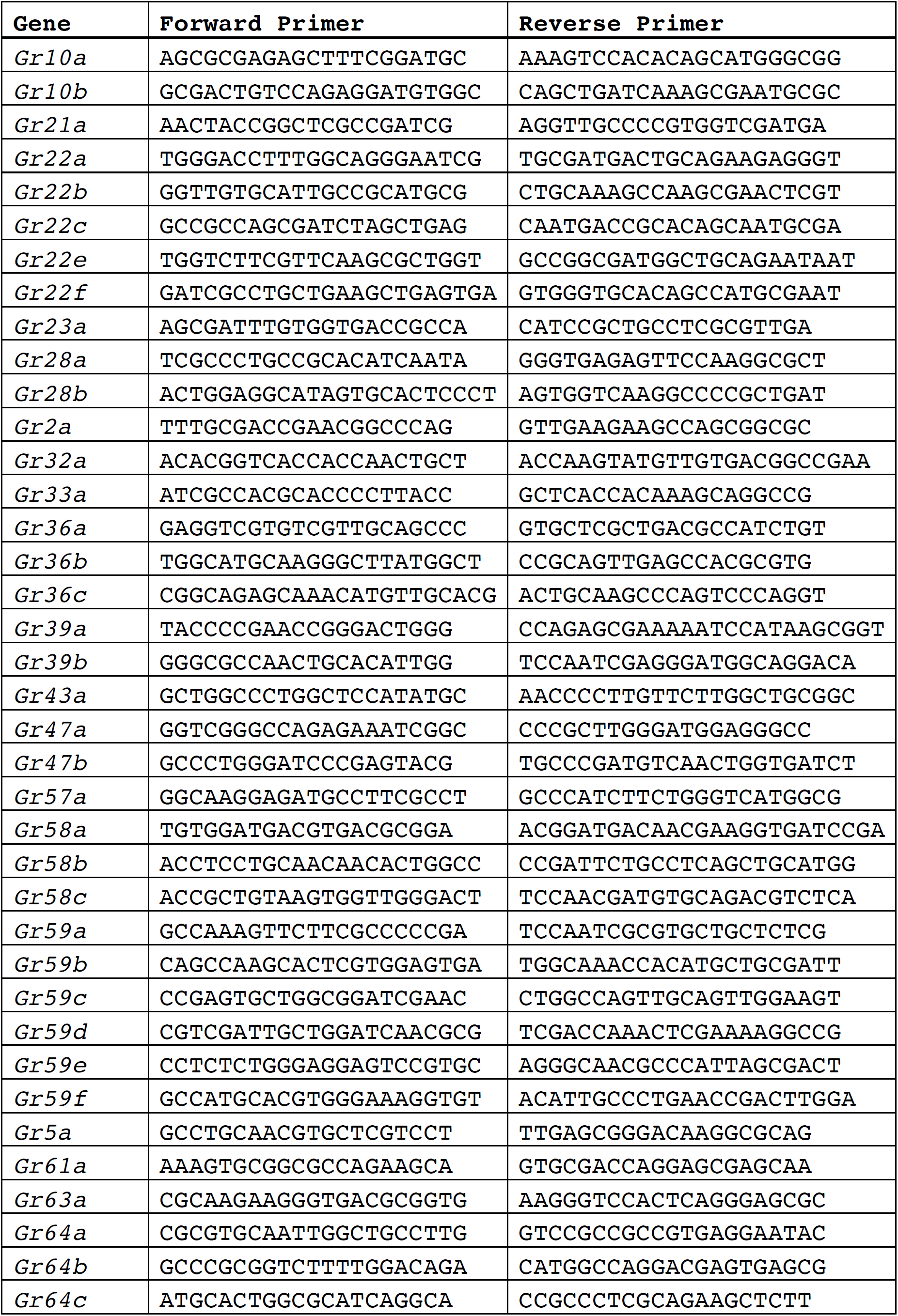
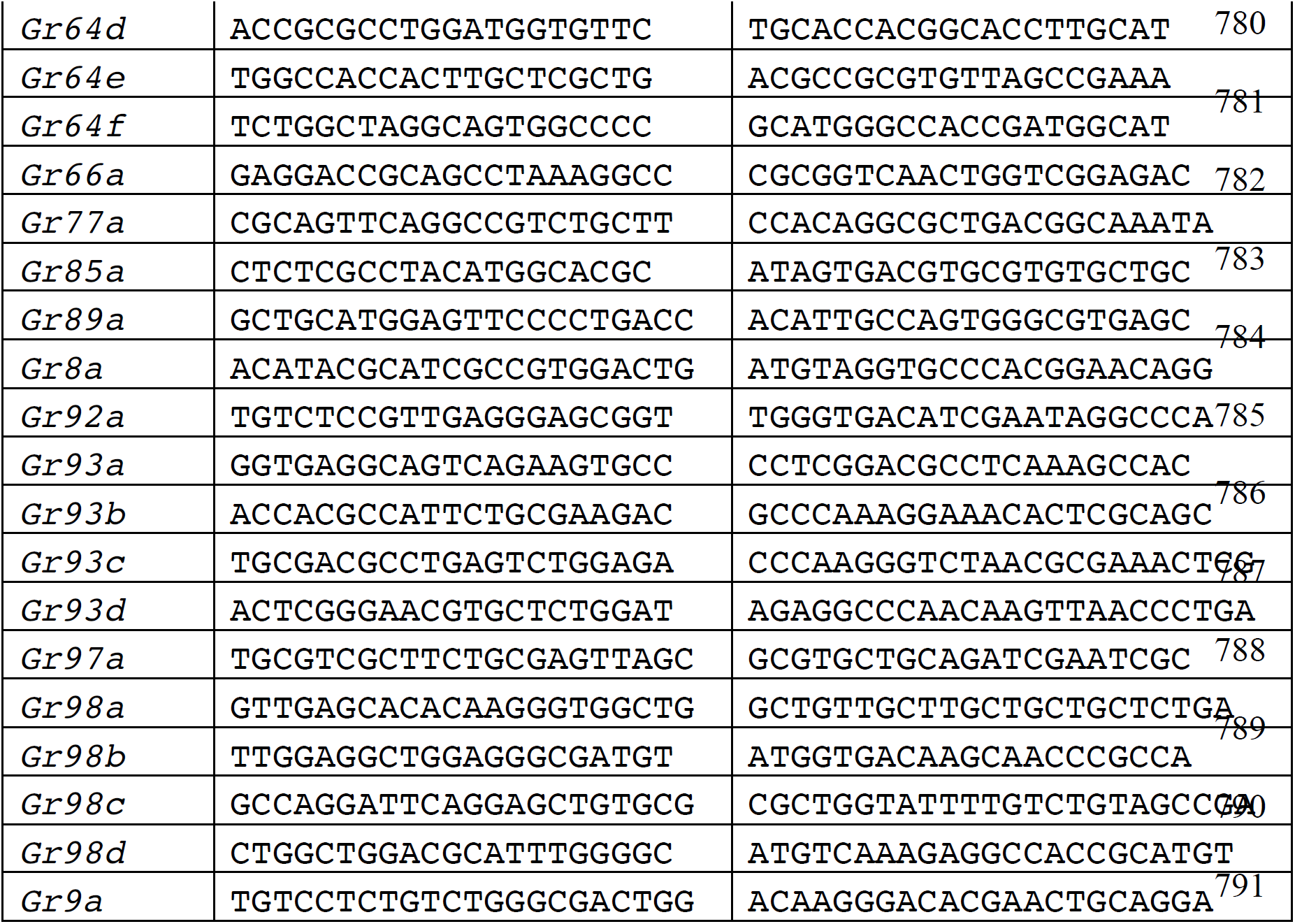
Nucleotide sequences for qRT-PCR primers for *D. melanogaster Gr* genes.

**Table 6.**
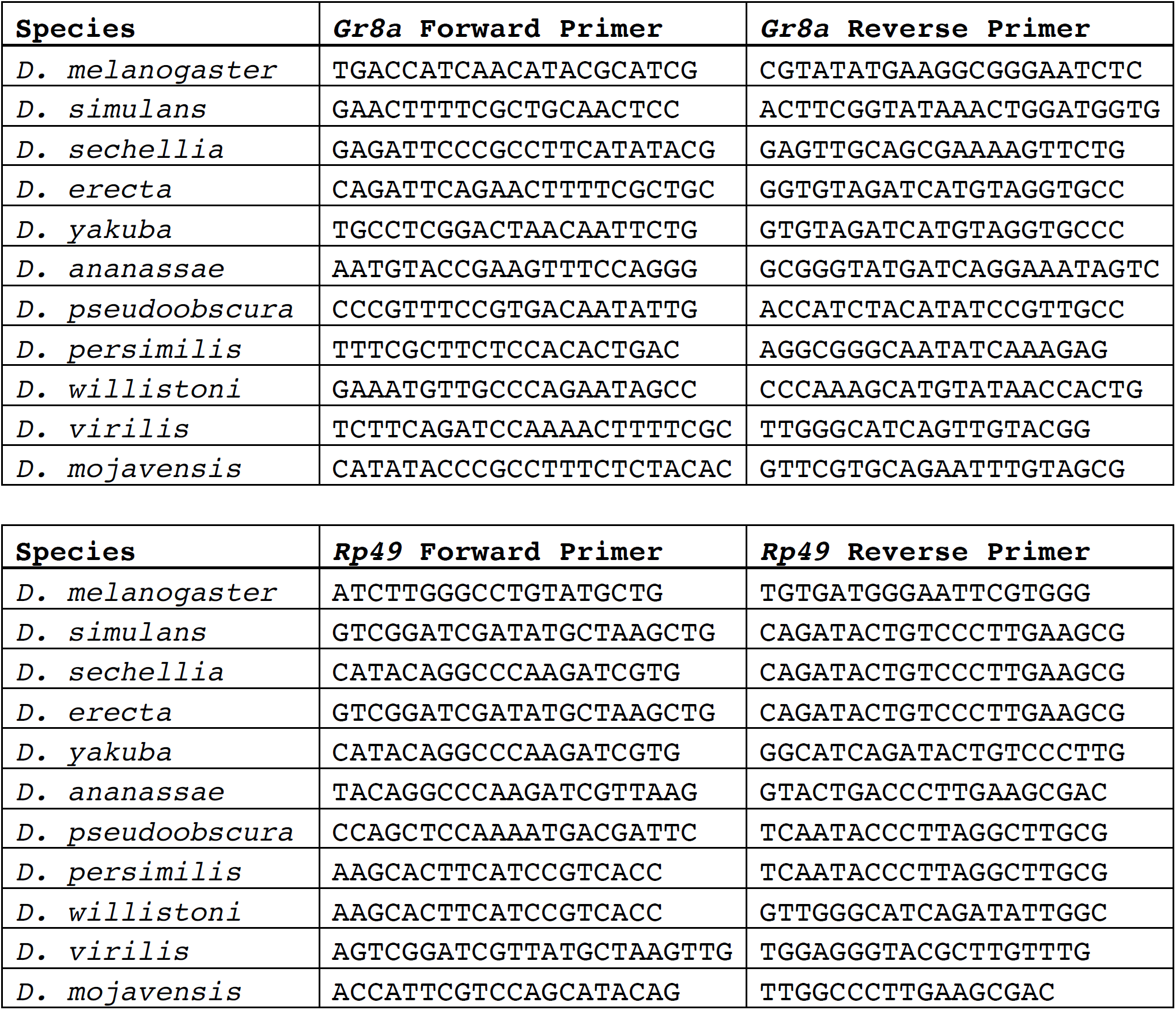
Nucleotide sequences for qRT-PCR primers for *D. melanogaster Gr8a, Rp49* and orthologs.

**Table 7.**
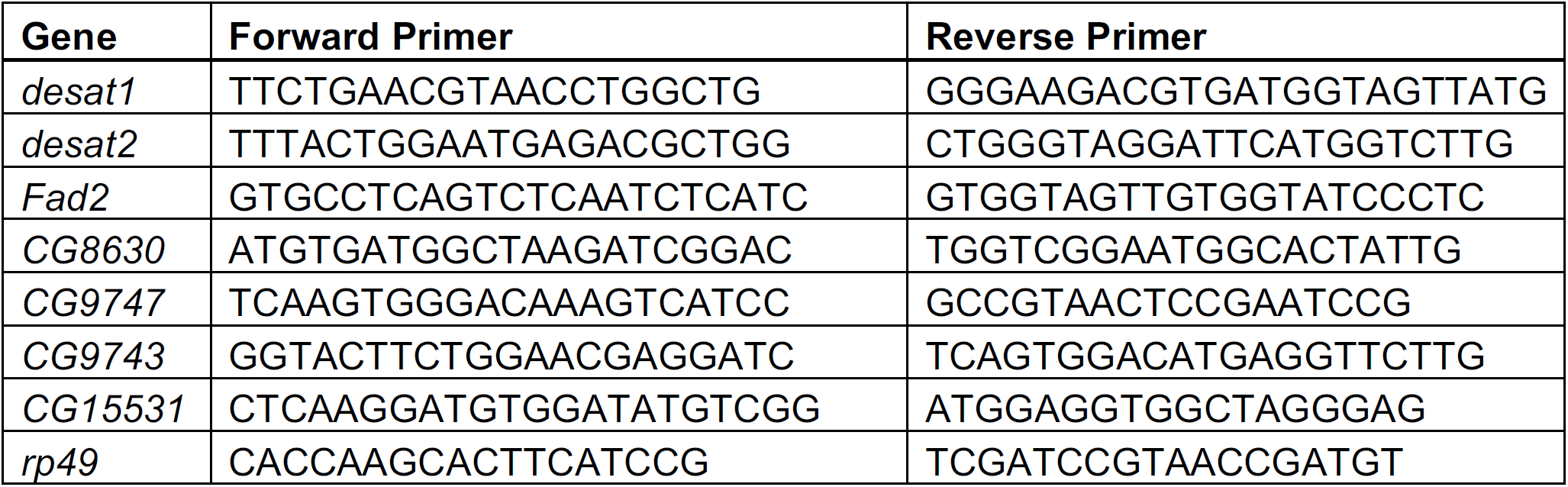
Nucleotide sequences for qRT-PCR primers for *D. melanogaster* desaturase enzyme genes.

### Courtship Behavior

Single-pair assays were performed as we have previously published (Lu et al., 2014, 2012). In short, newly eclosed males were kept individually on standard fly food in plastic vials (12 x 75mm). Newly eclosed virgin females were kept in groups of 10 flies. All behaviors were done with 4-7 day-old animals, which were housed under constant conditions of 25° C and a 12h:12h light-dark cycle. Courtship was video recorded for 10 min for male courtship and 15 min for female mating receptivity. Male courtship latency and index were measured as previously described (Lu et al., 2014, 2012). Female receptivity index was defined as the time from the initiation of male courtship until copulation was observed. Unless otherwise indicated, assays were performed under red light conditions.

### Fitness measures

Three-day old virgin male-female pairs were housed together for 24 hours after which males were discarded. Subsequently, groups of five females were allowed to lay eggs daily on fresh agar grape plates for five consecutive days. Fitness was measured as total number of eggs laid.

### Perfuming studies

Synthetic compounds were synthesized by J.G.M. Perfuming studies were performed using a modified protocol from (Billeter et al., 2009). In short, 3 mg of each compound was dissolved in 6 mL hexane (Sigma-Aldrich #139386-500ML) and 0.5 mL was pipetted into individual 2 mL glass vials fitted with 9mm PTFE lined caps (Agilent Crosslab, Santa Clara, CA, USA). The hexane was evaporated under a nitrogen gas flow, such that a residue of the compound was left around the bottom one-third of the vial. Control vials were prepared using hexane without a spiked compound. Vials were kept at -20°C until use. Flies used in these trials were collected as described above, kept in single sex groups and aged for 4 days on standard cornmeal medium at 25°C. 24 hours before perfuming, 20 flies of one or the other sex were placed in glass vials containing standard cornmeal medium (12 × 75mm). To perfume the flies, these groups of 20 flies were dumped without anesthesia into each 2 mL vial containing the compound of interest, and were vortexed at medium-low speed for 3 pulses of 20 seconds punctuated by 20 second rest periods. Flies were transferred to new food vials and were allowed to recover for one hour. Perfumed flies were then used in courtship behavior assays as described above and the remaining flies were used in pheromone analyses to verify compound transfer. The genotype of flies that were perfumed differed based upon the genotype with the lower amount of each compound as determined in Figure 3 (B, C, F). In all cases, compound transfer was verified by CHC extraction and GC/MS (Table 8).

**Table 8.** Amount (ng) of each perfumed compound measured in each sample of perfumed flies (5 flies per sample).

| Compound | Genotype | Sex | ng/sample | ng/fly |
| --- | --- | --- | --- | --- |
| C<sub>25</sub> | wt | male | 1350 | 270 |
| C<sub>25</sub> | wt | male | 1170 | 234 |
| C<sub>25</sub> | wt | male | 27730 | 5546 |
| 9-C<sub>25</sub> | Gr8a- | male | 67390 | 13478 |
| 9-C<sub>25</sub> | Gr8a- | male | 59380 | 11876 |
| 9-C<sub>25</sub> | Gr8a- | male | 1000 | 200 |
| 7-C<sub>25</sub> | Gr8a- | male | 43790 | 8758 |
| 7-C<sub>25</sub> | Gr8a- | male | 31330 | 6266 |
| 7-C<sub>25</sub> | Gr8a- | male | 24130 | 4826 |
| 5-C<sub>25</sub> | Gr8a- | male | 12050 | 2410 |
| 5-C<sub>25</sub> | Gr8a- | male | 14160 | 2832 |
| 5-C<sub>25</sub> | Gr8a- | male | 1870 | 374 |
| C<sub>27</sub> | Gr8a- | male | 890 | 178 |
| C<sub>27</sub> | Gr8a- | male | 720 | 144 |
| C<sub>27</sub> | Gr8a- | male | 263 | 52.6 |
| 7-C<sub>27</sub> | Gr8a- | male | 36710 | 7342 |
| 7-C<sub>27</sub> | Gr8a- | male | 21250 | 4250 |
| 7-C<sub>27</sub> | Gr8a- | male | 15910 | 3182 |
| C<sub>29</sub> | wt | male | 260 | 52 |
| C<sub>29</sub> | wt | male | 340 | 68 |
| C<sub>29</sub> | wt | male | 890 | 178 |
| 9-C<sub>25</sub> | wt | female | 14060 | 2812 |
| 9-C<sub>25</sub> | wt | female | 13830 | 2766 |
| 7-C<sub>25</sub> | wt | female | 23200 | 4640 |
| 7-C<sub>25</sub> | wt | female | 12780 | 2556 |
| 7-C<sub>27</sub> | wt | female | 1010 | 202 |
| 7-C<sub>27</sub> | wt | female | 1080 | 216 |

### Phylogenetic analysis

Protein sequences of GR8A orthologs from the 12 sequenced *Drosophila* reference genomes were aligned by using the ClustalW algorithm in the Omega package (Sievers et al., 2011), followed by ProtTest (v2.4) to determine the best model of protein evolution (Abascal et al., 2005). Subsequently, Akaike and Bayesian information criterion scores were used to select the appropriate substitution matrix. We then used a maximum likelihood approach and rapid bootstrapping within RAxML v 7.2.8 Black Box on the Cipres web portal to make a phylogenetic tree (Miller et al., 2010). Visualizations of the bipartition files were made using FigTree v1.3.1 (http://tree.bio.ed.ac.uk/software/figtree/).

### Pheromone Analysis

Virgin flies were collected upon eclosion under a light CO^2^ anesthesia and kept in single-sex vials in groups of 10 with 6 biological replications for each genotype and sex. Virgin flies were aged for 5 days on standard cornmeal medium at 25°C. To collect mated flies, both females and males were aged for 3 days before single mating pairs were placed in a standard fly vial with standard cornmeal food for 24 hours. The pair was then separated for 24 hours before collection. Copulation was confirmed by the presence of larvae in the vials of mated females several days later. On the morning of day 5, flies were anesthetized under light CO^2^ and groups of five flies were placed in individual scintillation vials (VWR 74504-20). To extract CHCs, each group of flies was covered by 100 uL hexane (Sigma-Aldrich #139386-500ML) containing 50µg/mL hexacosane (Sigma-Aldrich #241687-5G) and was washed for ten minutes. Subsequently, hexane washes were transferred into a new 2 ml glass vial containing a 350 uL insert (Thermo Scientific C4000-LV-1W) and were stored at -20°C until shipment to the Millar laboratory.

Analyses of CHC profiles were done by gas chromatography and mass spectroscopy (GC-MS) in the Millar laboratory at UC Riverside as previously described (Chung et al., 2014). Peak areas were measured, and data was normalized to known quantity of internal standard hexacosane (Sigma-Aldrich #241687-5G). The relative proportion of each compound in each sample was calculated and used in further statistical analysis. Qualitative data were analyzed through a permutation MANOVA using the ADONIS function in the vegan package of R with Bray-Curtis dissimilarity measures (Oksanen et al., 2017). Data were visualized using non-metric multidimensional scaling (vegdist function in the vegan package of R followed by cmdscale function in the stats package (Oksanen et al., 2017)) using Bray-Curtis dissimilarity, and either 2 or 3 dimensions in order to minimize stress to < 0.1. Quantitative data were analyzed by using a t-test or Mann-Whitney Rank Sum Test in R 3.3.2 (R Core Team, 2016).

## Data availability

All relevant data are available from the corresponding author upon request.

## AUTHOR CONTRIBUTIONS

K.M.Z., C.V., J.G.M. and Y.B-S designed experiments. K.M.Z., C.V., N.L., X.L., S.H., J.G.M. and Y.B-S collected and analyzed data. K.M.Z., C.V. and Y.B-S wrote the manuscript.

## ACKNOWLEDGMENTS

We thank members of the Ben-Shahar lab for comments on earlier versions of the manuscript. We thank the Millar laboratory (UC Riverside) for performing the CHC analyses. We thank Joshua Krupp for assistance with perfuming studies, Nabeel Chowdhury and Deanna Simon for assistance with qRT-PCR analysis, and Paula Kiefel for technical help with generating transgenic flies. This work was supported by NSF grants 1545778 and 1754264, and NIH grant NS089834 awarded to Y.B. Stocks obtained from the Bloomington Drosophila Stock Center (NIH P40OD018537) were used in this study. Wild type *Drosophila* species were obtained from the National *Drosophila* Species Stock Center at Cornell University.

## Figures and figure legends

**Figure 2—figure supplement 1.**
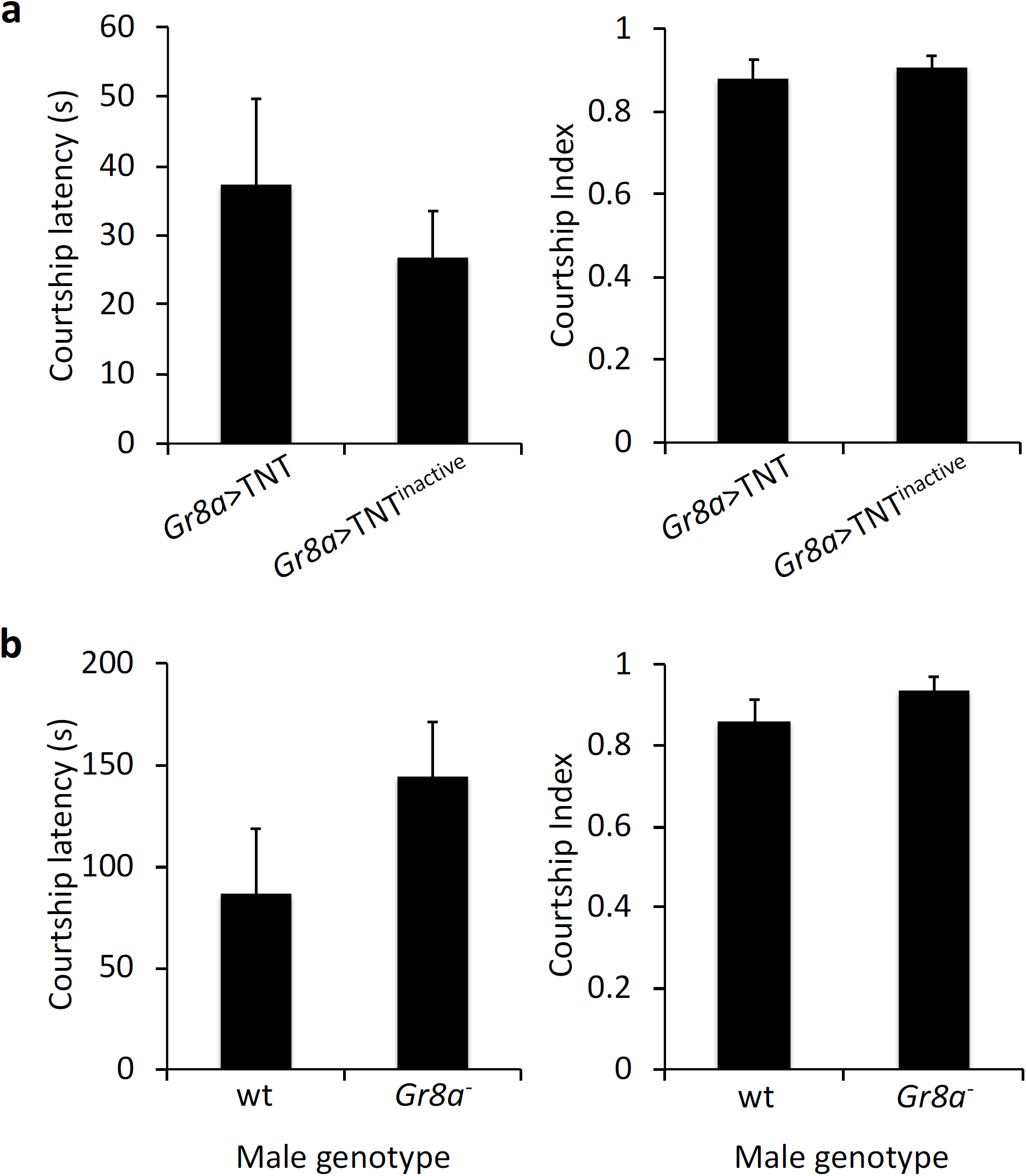
*Gr8a* has no effect on male Courtship Index or Latency toward wild-type females. (A) Courtship Index and Latency (s) of control *Gr8a-gal4/UAS-IMP-TNT- V1A* (*TNT*^*inactive*^) and *Gr8a-gal4/UAS-TNT-E* mutant males towards wild-type females. (B) Courtship index and latency (s) of wild-type (*CS*) and *Gr8a null* males toward wild-type decapitated females. Mann Whitney Rank Sum Test, not significant (p>0.05), n=15/group.

